# Motivation and Cognitive Control in Depression

**DOI:** 10.1101/500561

**Authors:** Ivan Grahek, Amitai Shenhav, Sebastian Musslick, Ruth M. Krebs, Ernst H.W. Koster

## Abstract

Depression is linked to deficits in cognitive control and a host of other cognitive impairments arise as a consequence of these deficits. Despite of their important role in depression, there are no mechanistic models of cognitive control deficits in depression. In this paper we propose how these deficits can emerge from the interaction between motivational and cognitive processes. We review depression-related impairments in key components of motivation along with new cognitive neuroscience models that focus on the role of motivation in the decision-making about cognitive control allocation. Based on this review we propose a unifying framework which connects motivational and cognitive control deficits in depression. This framework is rooted in computational models of cognitive control and offers a mechanistic understanding of cognitive control deficits in depression.

## Introduction

Depression^1^ profoundly influences the way in which we process information and think about ourselves, others, and the world around us. An individual suffering from depression will take a longer time to disengage from the processing of negative information and will experience difficulties in suppressing irrelevant thoughts, or shifting attention from one task to another in order to reach a goal. Such issues will make it difficult for that individual to regulate emotions and adapt to the changing environment. This is why cognitive processes are a crucial target for understanding and treating depression (Clark and Beck, 2010; Kaser et al., 2017).

Depression is characterized by impairments in attention, memory, and cognitive control (Millan et al., 2012). Cognitive control deficits are related to central features of depression such as concentration and memory problems and a host of other cognitive impairments and biases arise as a consequence of these deficits (Disner et al., 2011; Gotlib and Joormann, 2010). Cognitive control is crucial in motivated, goal-directed behavior. It represents a set of processes that allow for the flexible adaptation of cognition and behavior in accordance with our current goals (Botvinick and Cohen, 2014; Friedman and Miyake, 2017; Shenhav et al., 2013). For example, cognitive control is necessary if we want to inhibit negative thoughts and shift our attention to a new task. Impairments in such control processes are found across a wide range of psychiatric disorders (Millan et al., 2012), and they have been consistently linked with depressive symptoms (Snyder, 2013). However, our understanding of cognitive control in depression is limited in several important ways.

Current understanding of cognitive control in depression is predominately descriptive and research is focused on detecting deficits in specific cognitive processes, such as the inhibition of negative material (for a review see: Grahek et al., 2018). In spite of the important progress in charting cognitive control deficits related to depressive symptoms, the origin of these deficits remains poorly understood. It is not known why deficits in cognitive control develop, or how they are maintained. Most of the existing models view cognitive control deficits in depression as the reduced ability to exert control and do not offer mechanisms through which these deficits emerge. Currently there is a strong need for the development of a more mechanistic account of cognitive control deficits in depression. A mechanistic account moves beyond identification and description of a phenomenon. It does so by appealing to a mechanism: a structure defined by its components, their organization and interactions, which produce a phenomenon (Bechtel and Abrahamsen, 2005; Machamer et al., 2000). In this paper we argue that recent developments in research on motivation and cognitive control, as well as the development of computational theories of cognitive control, can contribute to such a mechanistic understanding of control deficits in depression. The use of such theories to explain cognitive control deficits in depression, instead of developing new models specific for psychopathology, holds the promise of advancing the understanding of cognitive control in both healthy and depressed individuals.

In this paper we first review disparate literatures on motivation and cognitive control in depression. Further, we describe computational models of cognitive control and demonstrate how they can be used to link motivation and cognition in depression. On the basis of this review, we rely on a computational model of cognitive control to propose a framework in which cognitive control deficits in depression arise from alterations in crucial components of motivation: reward anticipation, effort costs, and estimates of environment controllability. This view offers the possibility for re-conceptualizing depression-related cognitive control deficits. Instead of a *reduced ability* to employ control, we propose that control deficits can be viewed as changes in the decision-making process underlying cognitive control allocation. This decision-making process relies on crucial components of motivation: reward anticipation, effort, and estimates of the ability to control the environment. We use simulations of two cognitive tasks to demonstrate how this framework can be used to derive behavioral predictions about the impact of motivational impairments on cognitive control.

## Cognitive Control in Depression

Deficits in cognitive control have not only been documented in clinically depressed individuals (Snyder, 2013), but also in patients in remission (Demeyer et al., 2012; Levens and Gotlib, 2015), and in at-risk populations (Derakshan et al., 2009; Owens et al., 2012). Meta-analytic evidence from behavioral studies suggests that depression is reliably linked to deficits in cognitive control (Rock et al., 2014; Snyder, 2013). There is also emerging evidence that cognitive remediation training aimed at improving cognitive control processes reduces depressive symptoms (for a review see: Koster, Hoorelbeke, Onraedt, Owens, & Derakshan, 2017). At the neural level, depressive symptoms have been linked to changes in the activity of the dorsolateral prefrontal cortex (dlPFC) and anterior cingulate cortex (ACC; Davidson, Pizzagalli, Nitschke, & Putnam, 2002; Gotlib & Hamilton, 2008; Pizzagalli, 2011). Meta-analyses of the neuroimaging studies also point to the differences between healthy and depressed individuals, both in activation in these two regions during cognitive tasks (McTeague et al., 2017), as well as in the gray matter volume (Goodkind et al., 2015). However, the neuroimaging studies have often been conducted on very small samples, and there is considerable heterogeneity in their results (e.g., Müller et al., 2016; for a discussion see: Barch and Pagliaccio, 2017). Multiple authors have proposed that the reduced activity in the dlPFC and the ACC is related to the diminished ability of depressed individuals to employ cognitive control (Disner et al., 2011; Joormann, Yoon, & Zetsche, 2007).

Cognitive control processes are considered to be an important vulnerability factor for depression. Cognitive impairments in attention, interpretation, and memory may arise as a consequence of control deficits (Gotlib and Joormann, 2010; Millan et al., 2012; Siegle et al., 2007). For example, Gotlib and Joorman (2010) have suggested that depressed individuals’ difficulties in disengaging attention from negative stimuli, or forgetting such stimuli, could be caused by cognitive control deficits. This proposal has recently received empirical support (Everaert et al., 2017). Lowered levels of cognitive control increase and sustain depressive symptoms via their proximal links with emotion regulation strategies such as rumination (Joormann & Vanderlind, 2014; Whitmer & Gotlib, 2013). Research on cognitive control in depression has been focused on charting deficits in different cognitive control processes. Specific deficits in processes such as inhibition, shifting, and updating have been documented (Joormann & Tanovic, 2015). These deficits are commonly thought of as the *lowered ability* to inhibit certain thoughts or stimuli, shift attention away from them, or update the contents of working memory. However, it is important to note that not all accounts of cognitive impairments in depression postulate a reduced ability. For example, the cognitive-initiative account of memory in depression focuses on changes in initiative – a concept close to motivation – to explain memory impairments in depression (Hertel, 2000, 1994).

In a recent analysis of theoretical models of cognitive control in depression (Grahek et al., 2018) we identified three main conceptual problems in the field: (1) the use of descriptive models of cognitive control, (2) the reliance on describing the impairments instead of searching for mechanisms, and (3) the lack of integration between cognitive, motivational, and emotional impairments. These issues are hindering further progress in understanding how and why cognitive control is impaired in depression. In order to overcome some of the problems that we have outlined in our earlier work, in this paper we propose an integrated framework that links alterations in motivational processes with cognitive control deficits. This framework allows us to move away from the view that cognitive control deficits in depression stem from a reduced ability to exert control. Instead, we will argue for a view in which the deficits arise as a result of altered expectations about the value of exerting control.

## Components of Motivation in Depression

Cognitive impairments in depression are closely linked to impairments in emotional and motivational processes (Crocker et al., 2013). A wealth of depression research has focused on the relationship between cognitive and emotional processes. This approach has led to insights in key deficits related to the disengagement from emotionally negative material (Koster et al., 2011) and the ability to deploy cognitive control over emotional material (Joormann, 2010). While the processing of negative material and the presence of negative affect have been studied in relation to cognitive control impairments in depression, motivational deficits remain largely unexplored in this context. This is why we focus this paper on the link between motivation and cognition in depression. The links between motivation and cognitive processing of emotional material our out of the scope of the current paper.

Motivation is goal-directed when effort is invested in order to bring about desired outcomes (Braver et al., 2014). Here we will focus on components of motivation that are relevant for goal-directed behavior because of their relevance for cognitive control processes. Motivated goal-directed behavior is flexible and sensitive to the current state of the individual and the environment. Two types of representations are crucial in driving this type of behavior: 1) action-outcome contingencies, and 2) the value of potential outcomes (Balleine and O’Doherty, 2010; Dickinson, 1985; Wood and Rünger, 2016). Action-outcome contingencies represent the probability that an action will result in a desired outcome. We will refer to these representations as *outcome controllability* (one’s estimate of their ability to control outcomes in an environment) and *outcome value* (the expected reinforcement - total reward and/or punishment - for reaching an outcome). The third concept that we will consider is effort, a variable that is central to the study of motivation. Effort represents the intensification of physical or mental activity needed to reach a goal (Inzlicht et al., 2018; Kurzban et al., 2013). This intensification comes with a cost, and we will refer to the effort requirements for reaching an outcome as *effort costs.*

Goal-directed motivated behavior emerges with the integration of these three components. For example, imagine a person working long hours in order to get a promotion at work. This behavior is motivated and goal-directed because this person believes that working hard (high effort) will lead to the promotion (high outcome controllability), and the promotion is desired (high outcome value). As we review next, there is evidence that each of these three components of motivation can be impaired in depression (Barch et al., 2015; Griffiths et al., 2014).

### Outcome Controllability

The classic paradigm for investigating the role of outcome controllability in response to stressors was developed by Seligman and Maier (1967). They demonstrated that animals who were subjected to uncontrollable stressors (inescapable shocks) exhibited passivity, a phenomenon they referred to as learned helplessness. They found that controllable stressors (escapable shocks) did not induce learned helplessness. Overall, uncontrollable stressors were found to induce responses that resemble some of the symptoms of depression (Maier, 1984; Maier and Watkins, 2005). Further work by Maier and colleagues (2006) revealed that animals were able to detect the possibility of control over their environment. While animals have a default reaction of passivity when experiencing stress, this default response can be overcome by learning that the stressors are controllable. The medial prefrontal cortex (mPFC) in rodents detects whether a stimulus is controllable and inhibits the default passivity in response to shocks. In this way, the ability to exert control over the environment serves as a protective factor against negative behavioral and physiological effects of stress (Maier and Seligman, 2016). Seligman and colleagues demonstrated that learned helplessness is also observed in humans, and suggested that it has significant relevance for understanding depression and related disorders (Abramson, Seligman, & Teasdale, 1978; Maier & Seligman, 2016; Seligman, 1972).

The idea that environmental controllability is crucial for one’s health, security, and well-being is supported by several other lines of research (Leotti et al., 2010). For instance, Moscarello and Hartley (2017) have recently proposed that goal-directed behavior is strongly influenced by estimates of agency. The authors propose that animals and humans infer their ability to control their environment by ascertaining the relationship between their actions and motivationally significant outcomes. These estimates are generalized and determine the probability of being able to exert control in a novel environment. Estimates of agency are used to calibrate ongoing behaviors. If the estimates of agency are high (i.e., high probability of being able to control the environment), goal-directed behavior is promoted. If the estimates are low, behavior is more likely to rely on an innate reactive repertoire. In this way, controllability of the environment, inferred from previous learning, is crucial for promoting either goal-directed or habitual behavior (Liljeholm, Tricomi, O’Doherty, & Balleine, 2011; Miller, Shenhav, & Ludvig, in press). Lowered estimates of outcome controllability, resulting in alterations of goal-directed behavior, could be an important factor in depression.

Recent theoretical frameworks of depression also emphasize the importance of outcome controllability. For example, Pizzagalli (2014) has proposed a model of anhedonia in depression in which stress plays a central role in the development of anhedonic symptoms. According to Pizzagalli, certain stressors, especially if they are uncontrollable, induce anhedonic behavior by causing dysfunctions in mesolimbic dopamine pathways crucial for motivated behavior. De Raedt and Hooley (2016) have argued that individual’s expectancies about their ability to cope with future negative events play a crucial role in depression. These expectancies are proposed to be formed based on previous coping experiences and are one factor that determines the actual ability to cope with stressors when they occur. Notably, these authors suggest that expectancies influence the ability to cope with stressors by modulating the proactive allocation of cognitive control prior to encountering a stressful situation. In sum, altered estimates of outcome controllability seem to be an important factor contributing to the levels of depressive symptoms.

### Outcome Value

Representations of outcome value are also strongly altered in depression. In recent years, motivational impairments in depression have received significant attention and a more fine-grained view of these impairments is emerging (Pizzagalli, 2014; Treadway and Zald, 2013). Anhedonia is one of the two core symptoms of depression and is defined as a loss of pleasure in previously enjoyable activities or a loss of interest in pursuing them (DSM-5; American Psychiatric Association, 2013). Anhedonia is a good predictor of antidepressant treatment success (Uher et al., 2012), the course of depression (Spijker et al., 2001; Wardenaar et al., 2012), and the time to remission and number of depression-free days after antidepressant treatment (McMakin et al., 2012). In spite of the importance of anhedonia in depression, the conceptualization and measurement of anhedonia has been heterogeneous and inconsistent (for a discussion see: Rizvi, Pizzagalli, Sproule, & Kennedy, 2016). Traditionally, anhedonia has been primarily viewed as an impairment in consummatory pleasure (i.e. liking rewards when they are obtained). However, the evidence for impairments in consummatory pleasure in depression is mixed (Barch et al., 2015; Treadway and Zald, 2013).

Animal models suggest that the mesolimbic dopamine system is selectively involved in reward motivation, but not in hedonic responses when rewards are gained (Haber and Knutson, 2010; Salamone and Correa, 2012). These insights have stimulated research on anhedonia in depression and schizophrenia, revealing impairments in motivation that are not related to hedonic responses. As a result, an emerging, more nuanced account of motivation in anhedonia, emphasizes multiple reward processing deficits such as anticipation of rewards, reinforcement learning, effort expenditure, and value-based decision making (Barch and Dowd, 2010; Der-Avakian and Markou, 2012; Pizzagalli, 2014; Romer Thomsen et al., 2015; Strauss and Gold, 2012; Zald and Treadway, 2017). These deficits point toward changes in outcome value. They influence how outcome values are learned, anticipated, and translated into behavior.

Depression is related to a number of reward processing deficits (for reviews see: Barch et al., 2015; Pizzagalli, 2014; Treadway & Zald, 2011). Converging evidence from self-report studies, behavioral tasks, physiological, and neuroimaging experiments suggests that depression, especially in the presence of anhedonia, is linked to reduced anticipation of rewards and impaired implicit reinforcement learning. For example, monetary incentive delay tasks (Knutson et al., 2000) were developed in order to decompose anticipatory (e.g., the period after notifying participants about a possible reward) and consummatory (e.g., receipt of a monetary reward) aspects of reward processing. In these tasks depressed individuals, relative to healthy controls, display reduced behavioral and neural responses in anticipation of rewards (Pizzagalli, 2014). The literature on motivational impairments in depression suggests that depressed individuals, especially in the presence of anhedonic symptoms, assign value outcomes in a manner that differs from healthy controls. This can be caused by anticipating lower payoffs and/or impaired reinforcement learning.

### Effort costs

Another important factor influencing motivated behavior is effort that needs to be expended in order to reach a desired outcome. Effort involves the expected cost necessary to reach an outcome. This cost is weighed against expected benefits in order to choose which actions to pursue (Wallis and Rushworth, 2014). Reduced effort exertion is associated with multiple mental disorders (Culbreth et al., 2018; Salamone et al., 2016). Clinical studies have focused on the exertion of physical effort in order to obtain rewards, demonstrating that anhedonia in depression is related to reduced effort exertion (for an excellent review see: Zald & Treadway, 2017). In order to investigate this process, Treadway and colleagues developed the Effort Expenditure for Rewards Task (EEfRT; Treadway, Bossaller, Shelton, & Zald, 2012), which involves having participants choose how much effort they want to expend in order to gain varying amounts of reward. Using this task, the authors have demonstrated that depressed individuals are less willing to exert effort than healthy controls (see also: Cléry-Melin et al., 2011; Yang et al., 2014). However, the research on effort exertion in anhedonia and depression has largely been focused on physical effort while ignoring cognitive effort.

Although cognitive effort has been an important topic of research for a long time (Kahneman, 1973), in recent years there has been an upsurge in cognitive and neuroscience research on this topic (Kool and Botvinick, 2018; Westbrook and Braver, 2015). More specifically, research on cognitive control has focused on the role of effort costs in decision-making about control allocation (for a recent review see: Shenhav et al., 2017). Cognitive control processes require more effort than automatic ones, and effort needs to be expended in order to override automatic processes in order to reach a goal. Research is starting to demonstrate that there are individual differences in the exertion of this type of effort (Westbrook et al., 2013). To date, there are not a lot of studies on cognitive effort in depression. However, a recent study has demonstrated the inverse relationship between depressive symptoms and a questionnaire-based measure of the willingness to exert cognitive effort (Marchetti et al., 2018). Also, the first experimental study of cognitive effort in depression revealed similar results to the results obtained with physical effort (Hershenberg et al., 2016).

## Cognitive Control as a Process Reliant on Motivation

Controllability, value, and effort of a desired outcome depend on previous learning. They have been studied extensively in the context of animal and human learning at both behavioral and neural levels (for an overivew see: Daw and O’Doherty, 2014). The importance of these components of motivated behavior has also been recognized in other fields of psychology and neuroscience. For example, in social psychology the concepts of self-efficacy (Bandura, 2001, 1977) and locus of control (Rotter, 1966) are closely linked to controllability. Also, concepts of feasibility and desirability of goals (Oettingen and Gollwitzer, 2001) correspond to the concepts of controllability and outcome value. Similar concepts can be found in psychology of motivation (Atkinson, 1957; Wigfield and Eccles, 2000) and work psychology (Bonner and Sprinkle, 2002). The role of motivation is now increasingly recognized in cognitive psychology and cognitive neuroscience.

Automatic or habitual responding can be suitable for a number of everyday situations, but for tasks that are more novel, uncertain, or complex, individuals need to engage cognitive control. This set of processes helps to overcome automatic response tendencies in favor of controlled modes of information processing and behavior. This allows for the coordination of our thoughts and actions in accordance with our goals. Recent research has started to emphasize the role of motivation in the timing, intensity, and direction of cognitive control allocation. It is now recognized that the allocation of cognitive control is driven by goals and therefore closely linked to motivation (Braver et al., 2014).

Reward prospect enhances cognitive control processes (for reviews see: Botvinick and Braver, 2015; Krebs and Woldorff, 2017). This effect has been observed on various tasks that tap into different components of control such as: attentional control (Padmala and Pessoa, 2011), response inhibition (Leotti and Wager, 2010), conflict adaptation (Braem et al., 2012), and task-switching (Aarts et al., 2010). The effect is not restricted to situations in which reward is signaled by advance cues augmenting preparatory control processes. Comparable results are found when reward is signaled simultaneously with the target thereby promoting fast control adjustments (Krebs et al., 2010). But why is there a need to adapt control in the first place? Is it not the most optimal strategy to always exert maximal control over our thoughts and actions? The emerging answer is that exerting cognitive control carries intrinsic effort-related costs (Shenhav et al., 2017). It has been shown that exerting control is effortful and that individuals tend to avoid it (Kool et al., 2010). Both reward prospect associated with an outcome, and the effort needed to reach that outcome are important factors in determining the way in which cognitive control is allocated.

Another important factor in determining how cognitive control is allocated is the learned contingency between actions and outcomes. Although this factor has received less empirical attention, several computational models of cognitive control emphasize its importance. For example, the model of Alexander and Brown (2011) stresses the importance of predicting mappings between responses and outcomes in cognitive control. The model proposed by Shenhav and colleagues (2013) posits that the allocation of cognitive control relies, among other factors, on the probability of an outcome conditional on the type and intensity of control. In this way, allocation of cognitive control depends on predictions about probabilities of certain actions (e.g., intense control allocation or a certain response) producing desired outcomes (e.g., solving a task correctly).

### Computational Models of Cognitive Control

The insight that cognitive control is closely related to motivation is formalized in a growing number of computational models of cognitive control. At present there are multiple theoretical and computational models which deal with the problem of how cognitive control is allocated (for recent reviews see: Botvinick & Cohen, 2014; Verguts, 2017). These models specify a set of components needed for optimal decision-making about control allocation and they are not mutually exclusive. Many of them include the representations of outcome value and/or outcome controllability (Brown and Alexander, 2017; Holroyd and McClure, 2015; Shenhav et al., 2013; Verguts et al., 2015). Also, many of these models assume that learning is crucial in forming these representations and emphasize the importance of learning for cognitive control (Abrahamse et al., 2016; Bhandari et al., 2017). Our framework relies on the current formulation of the Expected Value of Control (EVC) theory (Shenhav et al., 2013). This theory includes representations of outcome controllability, value, and cost in the decision-making process about cognitive control allocation.

The EVC theory provides a normative account of cognitive control allocation as resulting from a decision-making process in which potential gains of control allocation are weighted against their costs (Shenhav et al., 2013). The theory postulates that these decisions determine how control is allocated across candidate control signals. These control signals vary on two dimensions: *signal identity*, which determines the processes which are engaged (e.g., paying attention to the ink color in a Stroop task), and *signal intensity* which determines the amount of control allocated (e.g., how intensely to pay attention to the ink color). The theory proposes that control signals are selected in such a way that maximizes the expected value of control at any given moment.

The EVC for a given control signal within a given state is determined by three components: efficacy, value, and cost. *Efficacy* is defined as the probability of a certain outcome given a signal of a particular identity and intensity, and the current state. In other words, the efficacy component can be described as the probability that certain outcomes will occur (i.e., that our actions lead to what we desire) or simply as action-outcome contingencies inferred from previous experience. In standard cognitive control paradigms, such as the Stroop task, an outcome can be a correct or an incorrect response on a particular trial. In this context, we can assign a probability to each of the two possible outcomes given each of the possible signal intensities and identities. *Value* is assigned to each of the possible outcomes and it represents the value of an outcome in terms of possible rewards or punishments associated with the outcome. The rewards can be both intrinsic (i.e., motivation to do the task well) and extrinsic (i.e., monetary rewards for good performance). Punishments can also come from multiple sources such as monetary loss or poor task performance. Outcome values are also modulated by the time it takes for the outcomes to occur, as individuals try to maximize reward rates (rewards per unit of time) and not rewards per se. *Cost* is defined as the expected cost associated with the specified intensity of control allocation. Within the EVC theory, cost arises as the property of the neural system. This means that as the intensity of a signal to allocate control becomes stronger, so does the cost of allocating control. In sum, the EVC is determined by the probability of an outcome for a given control signal (efficacy), the value of that outcome, and the cost associated with control allocation.

At the neural level, the EVC theory was developed to explain the function of the dorsal anterior cingulate cortex (dACC). The theory assigns the critical role in decision-making about control allocation to this region. It is proposed that the dACC plays a key role in calculating the EVC and that the outcome of this process is then signaled to other regions which implement control, such as the dlPFC (Shenhav et al., 2016, 2013). The dACC is assigned a role in both monitoring for the changes in the environment relevant for control (rewards, punishments, errors, etc.), and, based on this, specifying which control signals maximize the EVC.

The EVC theory provides a detailed and formal account of how cognitive control is allocated. One of the main advantages of the theory is its ability to integrate a wide range of behavioral and neural findings. The computational implementation of the theory has been shown to account for various effects associated with the allocation of cognitive control such as the sequential adaptation effects and post-error slowing (Musslick et al., 2015), including how individuals can learn about features of their environment that predict incentives for and demands of control allocation (Lieder et al., 2018).

## Proposed Framework

We propose a framework that integrates disparate areas of research: cognitive control deficits and motivational impairments in depression. This framework encompasses the controllability, value, and effort of outcomes in order to provide a more mechanistic understanding of these deficits. Furthermore, the framework emphasizes the crucial role of learning in the process of allocating cognitive control. In this way the framework allows for the integration of motivational, learning, and cognitive control deficits in depression. We put forward direct links between components of motivation and reduced cognitive control in depression and propose how these links emerge through learning. In this way our framework offers a first step toward understanding how cognitive control deficits arise. In order to do so, we rely on the EVC theory which offers a computationally explicit model of cognitive control (Shenhav et al., 2013). This allows us to focus on the ways in which expected value of control might be reduced in depression, which results in allocating cognitive control with reduced intensity. We propose that cognitive control deficits in depression arise as a consequence of changes in the decision-making process about control allocation. These deficits in depression occur due to lowered expected value of control. The expected value of control relies on three components of motivation: controllability of outcomes, their value, and the effort needed to attain them.

We argue that research on motivation in depression provides insights into mechanisms leading to a reduction in the expected value of control. The efficacy component in the EVC theory is closely linked to what we have termed outcome controllability - the estimates of the probability that actions will lead to desired outcomes. These representations are central in guiding motivated behavior (Moscarello and Hartley, 2017). As we have reviewed, there is emerging evidence that estimates of outcome controllability and beliefs related to controllability of the environment, are changed in depression. The value component in the EVC theory is related to what we have termed outcome value - the expected reinforcers following an outcome. We have outlined how reward anticipation and reinforcement learning are impaired in depression, especially in the presence of anhedonic symptoms (Treadway and Zald, 2013). The cost component in the EVC theory is linked to the cost of effort needed to obtain rewards. We have reviewed how this process is altered in depression. All of these components of motivation rely on previous learning to estimate the probability of being able to exert control in a new environment, the value of outcomes in that environment, and the effort needed in order to reach them. These insights from the study of motivation in depression offer evidence for the reduction in efficacy, value, and cost of control. This leads to lower expected value of control which in turn results in reduced allocation of cognitive control (Figure 1).

**Figure 1.**
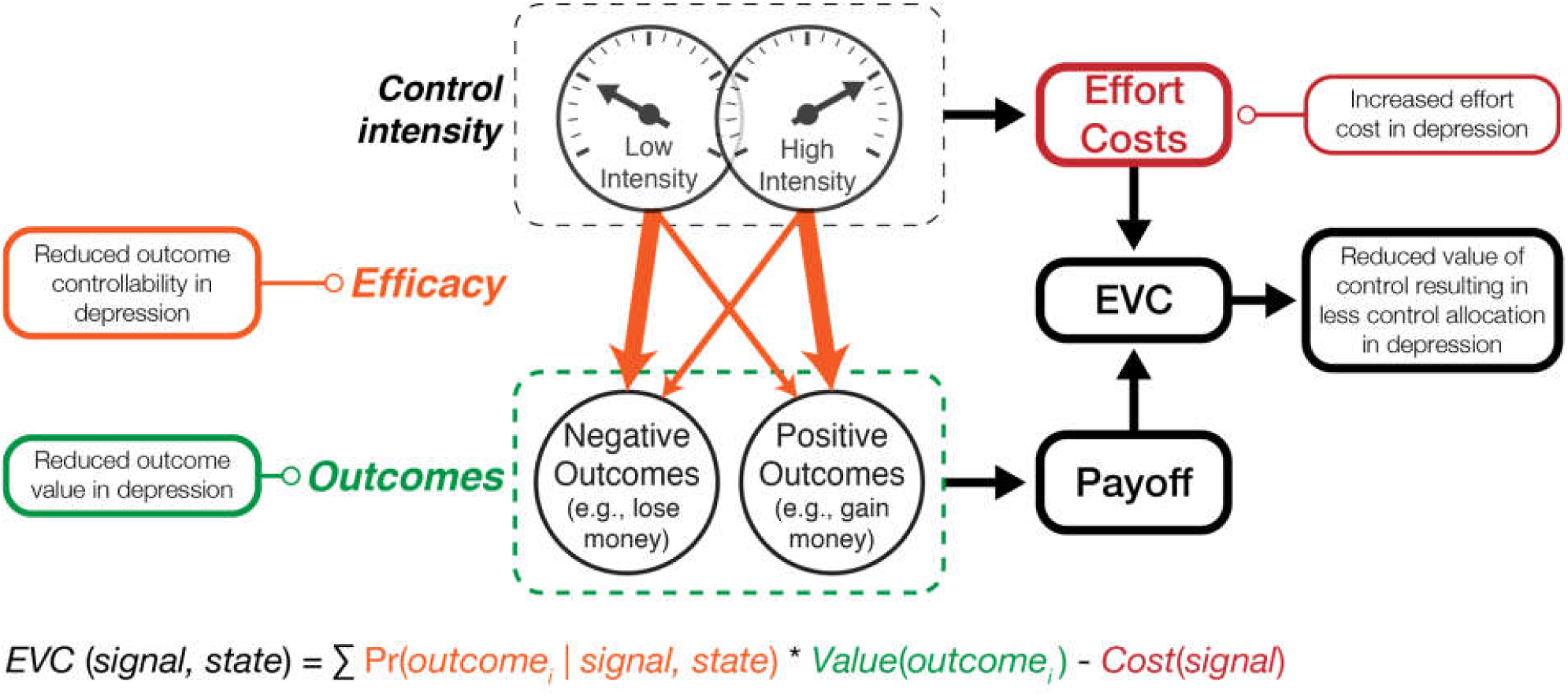
The schematic representation of the proposed framework. Depression-related changes in the outcome value (e.g. reduced reward anticipation), outcome controllability (e.g. lowered estimates of controllability), and effort costs (e.g. reduced effort exertion), lead to the reduced value of control. This leads to the lowered amounts of control being allocated. The expected value of control (*EVC*) for a signal of a given intensity is calculated as the sum of the values of each possible outcome weighted by the probability of reaching that outcome for the given signal. The cost of allocating control is subtracted from that sum. The figure was adapted from Shenhav et al., 2013 with permission from the authors.

Without a doubt, estimates of controllability, value, and effort of outcomes, are formed through prior learning. Estimates formed in one situation are generalized and used in novel similar environments. For example, imagine a person who developed low estimates of controllability in the work environment. Over the years that person may have learned that no matter what they do in the work environment, it is unlikely that desired outcomes occur. That person will tend to allocate less cognitive control in that environment and will develop beliefs about the inability to do the job well. This will lead to a reduction in motivated behavior and will further strengthen existing low controllability estimates and beliefs. Alternatively, another individual might estimate high controllability in the same work environment. However, that individual can anticipate low amounts of rewards associated with achieving good results in work, or high amounts of effort needed to do so. That individual does not anticipate pleasure in work anymore and will also tend to allocate less cognitive control during work hours.

Within our framework, depression-related deficits in cognitive control processes such as inhibition, task switching, or working memory updating (Joormann & Tanovic, 2015), can be conceptualized as products of changes in the expected value of control. In this way our framework integrates cognitive research on depression with impairments in other domains. Our framework outlines the links between cognitive control deficits and the study of motivation in depression. The framework can account for both the existing research findings and further develop the field by connecting these findings to previously unrelated literatures. Within our framework, cognitive control deficits in depression (Gotlib and Joormann, 2010) can be studied in relation with reward processing impairments in depression (Admon and Pizzagalli, 2015) and changes in perceived controllability of an environment (Moscarello and Hartley, 2017).

This framework has several key implications. First, cognitive control deficits do not have a single cause and can be caused by impairments in different processes. Deficits can thus be present in depressed individuals with different clusters of symptoms (e.g., mainly depressive mood or mainly anhedonia). Second, cognitive control deficits are causally related to reward processing impairments, effort costs, and estimates of controllability of an environment. These impairments influence allocation of cognitive control through learned estimates of reward probability, action-outcome contingencies, and needed effort. Third, cognitive control deficits are not a product of the simple reduced ability to exert control. They are a result of lowered expectations about the value of exerting cognitive control.

The proposed framework goes beyond the current understanding of cognitive control deficits in depression. It does so by providing a more mechanistic understanding of such deficits. First, it identifies the three crucial components that give rise to these deficits. Second, it proposes that the three components play a crucial role in the lowered expected value of control which, in turn, leads to deficits in cognitive control. Further, we propose below how this mechanistic understanding can be used to derive model-based behavioral predictions, as well how it can be related to the neurobiology of depression.

### Simulation-based Behavioral Predictions

The proposed framework outlines the three key motivational components which determine how cognitive control is allocated. By relying on a formalized model of cognitive control, the framework is able to generate precise predictions about the influence of each of these components on the behavioral performance in tasks which require cognitive control. In this section we describe how the computational implementation of the EVC theory can be used to make such predictions. This allows our framework to go beyond the currently existing data and make testable predictions that can be falsified or corroborated in future studies.

After the original formulation of the EVC theory (Shenhav et al., 2013), the more recent work has developed a computational implementation of the theory (Lieder et al., 2018; Musslick et al., 2015). In the computational implementation, performance of each task (e.g., responding to the color or to the word of a Stroop stimulus) is implemented as a process of evidence accumulation toward a decision boundary. Allocation of control can modify the parameters of this decision process (e.g. the rate of evidence accumulation) to improve performance, depending on the current goal and environment. This implementation relies on the drift-diffusion model of decision making, which has been widely used to describe decision-making (Forstmann et al., 2016; Ratcliff et al., 2016), value-based choice (Krajbich and Rangel, 2011; Tajima et al., 2016), as well as cognitive control processes (Bogacz et al., 2006; Cohen et al., 1990; Dunovan et al., 2015; Dutilh and Vandekerckhove, 2013; Schmitz and Voss, 2012; Ueltzhöffer et al., 2015).

The computational implementation of the EVC theory (Lieder et al., 2018; Musslick et al., 2015) has been successfully used to simulate a wide range of the existing empirical results in the domain of cognitive control (e.g. congruency effects, congruency sequence effects, task-switching costs) and the influence of motivation on cognitive control (incentive-based enhancements in the processes such as inhibition, task-switching, and conflict adaptation). Here we use this model in order to provide behavioral predictions about cognitive control in depression. Specifically, we explore how the depression-related changes in the efficacy, cost, and value components should influence behavior on tasks that require cognitive control. These simulations, based on a model that predicts well the existing data in healthy individuals, provides clear predictions related to cognitive control allocation in depression.

To demonstrate how control-demanding behavior is affected by changes in the decision-making process about control allocation, we simulated the behavior of an agent across two paradigms while varying parameters of that agent’s EVC-driven control valuation (Musslick et al., 2015; for a detailed description of these simulations, see Supplementary Materials). We first simulated performance in a Stroop task in which the simulated agent had to categorize a the ink color (e.g. red or green) of a color word (e.g. “RED” or “GREEN”) on each trial (Stroop, 1935). We assessed the overall control allocation and the resulting performance cost associated with responding to a color that is response-incongruent with the word (e.g. “RED” displayed in green) compared to responding to a color that is response-congruent with a word (e.g. “RED” displayed in red). We also simulated behavior in a cognitive effort discounting (COGED) task, in which the agent must choose between performing a difficult, high-demand task for $2 and an easy, low-demand (baseline) task for a variable amount of monetary reward on each trial. We assessed the subjective value of performing a high-demand task by measuring amount of monetary reward offered for the baseline task for which the EVC agent is indifferent between the two tasks, and by normalizing this amount by the reward offered for the high-demand task ($2). We evaluated how these complementary measures – of task performance vs. task preference – were influenced by changes in the agent’s (a) cost of cognitive control (b) sensitivity to reward, and (c) expected efficacy of exerting cognitive control (control efficacy).

Consistent with observations in healthy human subjects, our simulated agents demonstrated an incongruency cost, generating more errors when performing incongruent trials compared to congruent trials (Stroop, 1935). Critically, agents chose to exert less control (Fig 1A-C), resulting in higher incongruency costs (Fig 1D-F) when they had (i) higher costs of control, (ii) lower reward sensitivity, and/or (iii) when they perceived their control as being less efficacious. The simulated agent also replicated the behavior of healthy human subjects in studies of cognitive effort selection, assigning lower subjective values to tasks that demanded higher amounts of cognitive control (Westbrook et al., 2013). However, the subjective value for a given task decreased as (i) the cost of cognitive control increased, (ii) reward sensitivity decreased, and/or (iii) the control efficacy decreases (Figure 2G-I). Interestingly, the influence of these changes in parameters are slightly magnified for high task difficulties compared to low task difficulties.

**Figure 2.**
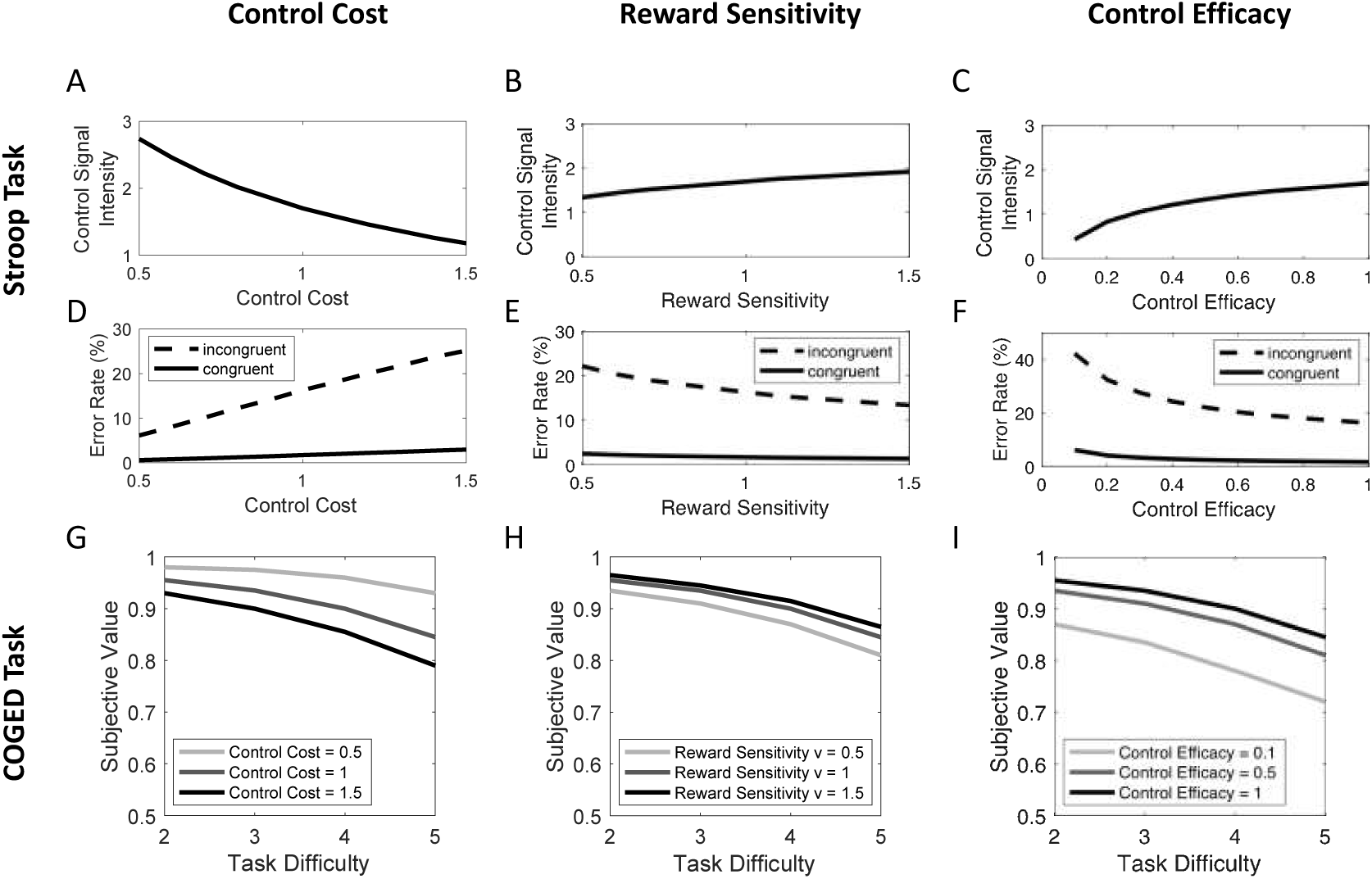
Effects of control cost, reward sensitivity and control efficacy on simulated behavior of the EVC model. Behavior of model was simulated in the Stroop paradigm and COGED paradigm (see supplementary Materials for details). (A-C) The amount of cognitive control allocated in a Stroop task is shown as a function of control cost, sensitivity to reward and expected control efficacy. (D-F) The error rate on incongruent and congruent trials is shown as a function of the three model parameters. (G-I) The subjective value of a task in the COGED paradigm is plotted as a function of task difficulty for different values of control costs, sensitivity to reward and control efficacy.

The computational implementation of our framework offers a possibility of studying cognitive control deficits in depression within the developing framework of computational psychiatry (Huys et al., 2016; Montague et al., 2012). Several computational frameworks have already started to model behavioral control in depression (Huys et al., 2015; Huys and Dayan, 2009) and our framework offers the possibility of extending this research into the domain of cognitive control deficits in depression. Our framework provides a clear path forward by applying a normative theory of cognitive control to understand cognitive control impairments in depression. Further, it points to the crucial components which interact to produce control deficits. We have described the advantages of the EVC theory above, but we do recognize the importance of other related computational models of cognitive control. Outcome value, outcome controllability, and effort costs are being recognized as crucial in allocation of cognitive control across different models of control. Because of this, our review of the relevant literature, and the proposed framework, will be useful in guiding future research beyond the boundaries of the EVC theory. The further use of the computational models developed in cognitive neuroscience to understand cognitive control deficits in depression will advance the field in several ways. First, it will allow for the more unified understanding of cognition in both healthy and in individuals suffering from mental illnesses. Second, it will avoid the creation of models that are tailored specifically to understand cognition in depression. Finally, the application of these models to depression, and other mental disorders, will be able to inform and help improve the existing computational models of cognitive control.

### Neural Level

Cognitive control deficits in depression are proposed to be related to functional changes in the dlPFC and the ACC (Disner et al., 2011). This view is based on the conceptualization of cognitive control functions as reliant purely on the prefrontal cortex. However, this view has been challenged by the discovery of the role of other regions, such as the basal ganglia and corticostriatal loops in cognitive control and goal-directed behavior (for reviews see: Balleine & O’Doherty, 2010; Chatham & Badre, 2015; Haber, 2016). By building on the EVC theory, our framework offers a more specific view on the role of these two regions in depression. We propose that depression-related changes in the decision-making process about control allocation are related to the activity of the dACC. These changes can further result in lower levels of dlPFC activation which implements control based on the signals from the dACC. In this way, depression is not related to a lowered ability of the dlPFC to implement control, but related to changes in inputs to the dACC used in decision-making about control allocation. In line with the importance of the dACC in our framework, current research has pointed to the crucial role of the ACC in depression (Holroyd and Umemoto, 2016; McTeague et al., 2017), as well as to the specific role of the dACC (Goodkind et al., 2015). Our framework also points to the important role of efficacy and value representations which serve as inputs in the decision-making process. In this way, it can guide further research in connecting the role of the dACC and other regions and networks related to encoding value, reward processing, and efficacy (Haber and Knutson, 2010; Moscarello and Hartley, 2017; Treadway and Zald, 2011).

### Relationship with Other Constructs Relevant for Depression

Lowered levels of cognitive control have a reciprocal relationship with negative beliefs and attributions. Negative beliefs about the self, the others, and the world represent a crucial cognitive vulnerability factor to depression (Beck, 1972). Attributions that are global, internal, and stable are also an important vulnerability factor (Abramson et al., 1989). Cognitive control deficits can strengthen those beliefs, but are also strengthened by them. Interestingly, there is already some progress on studying the maladaptive beliefs within the framework of computational psychiatry (Moutoussis et al., 2017). Finally, lowered levels of cognitive control are related to maladaptive use of emotion regulation strategies such as rumination (Nolen-Hoeksema et al., 2008) which contribute to the onset and maintenance of depressive symptoms (Joormann & Vanderlind, 2014). Impaired cognitive control also affects other cognitive processes thus producing cognitive biases in attention, interpretation, and memory, which further promote the maladaptive use of emotion regulation strategies (Everaert et al., 2012; Gotlib and Joormann, 2010).

An important research line has focused on the relationship between depressive symptoms and cognitive control over affectively negative material. These studies have demonstrated specific impairments in shifting attention away from negative material, removing negative material from the working memory, and inhibiting negative stimuli (Gotlib and Joormann, 2010; Joormann and Vanderlind, 2014; Koster et al., 2011). In its current form, our framework does not focus on these deficits. However, we hope to integrate this body of work in the framework in future studies. One of the interesting possibilities for such integration is the influence of negative material on efficacy estimates through prior learning.

## Future Directions and Open Questions

The proposed framework opens novel avenues for research, namely the links between cognitive control in depression and alterations in components of motivation. Future studies in this domain can more directly test some of the implications of our framework. To date there are not many studies that have investigated the links between components of motivation and cognitive control deficits in depression. We hope that our framework will inspire more studies in this direction. Here we outline some of the existing evidence and propose the paradigms that could be used in future research.

Several studies have already demonstrated the importance of motivation for cognitive functioning in depression (Moritz et al., 2017; Scheurich et al., 2008). Recently, studies have focused on more specific components of motivation. For example, several studies have demonstrated that reward-based improvements in cognitive control and attention are related to depressive symptoms (Anderson et al., 2014; Jazbec et al., 2005; Ravizza and Delgado, 2014). Future studies should focus on precise distinctions between different types of impairments related to reward processing. In this context, we believe that reward anticipation and effort costs are the most interesting processes to investigate in relation to cognitive processes in depression.

Depression research can rely on paradigms developed in cognitive science to study motivation and cognitive control (Botvinick and Braver, 2015). The study of effort expenditure is also gaining a great deal of attention and paradigms to study physical effort already exist (Treadway et al., 2012). For depression research, novel insights on cognitive effort (Shenhav et al., 2017) can be of particular relevance. Also, the relationship between anhedonic symptoms and cognitive control deficits in depression will be a crucial next step in better understanding cognition in depression. In this domain research should be guided by recent developments in measuring anhedonia through behavioral tasks and questionnaires (for a review on measures of anhedonia see: Rizvi et al., 2016).

In the domain of efficacy, there are no readily available paradigms that could directly inform us about its relationship with cognitive control. However, developments in the field of learned helplessness suggest that the interactions between stress and controllability are crucial for stress regulation (Maier and Seligman, 2016). Recent research is starting to provide paradigms to study stress controllability in humans (Bhanji et al., 2016; Hartley et al., 2014). The critical next step is to develop paradigms that would allow the investigation of cognitive control processes in relation to controllability. Recently, paradigms aimed to investigate different components of motivated action and learning have been developed in schizophrenia research. Using a novel paradigm, Morris and associates have demonstrated impairments in action-outcome learning in schizophrenia (Morris, Cyrzon, Green, Le Pelley, & Balleine, 2018; Morris, Quail, Griffiths, Green, & Balleine, 2015; see also: Liljeholm et al., 2011). The use of similar paradigms would allow for further work on computational models of motivation and cognition in depression. Recent computational work has demonstrated that generalization of learned action-outcome contingencies can account for a wide range of behavioral features associated with learned helplessness in animals and humans (Huys and Dayan, 2009; Lieder et al., 2013). However, there is a strong need for more empirical work which could precisely measure the importance of outcome controllability in depression.

Cognitive control deficits (as well as anhedonia) are transdiagnostic in nature and co-occur in many disorders (Goschke, 2014; Whitton et al., 2015). Although our model is focused on depression, the links between cognitive control and reward processing are relevant for other psychiatric disorders as well. Future studies in this domain hold the promise of not only detecting transdiagnostic constructs such as the positive valence and cognitive systems (Cuthbert and Insel, 2013; Insel et al., 2010), but starting to investigate the relations between these systems. In this context, the relationship between reward processing (positive valence system) and cognitive control (cognitive system) across different clinical populations can be of great interest.

In current research, depression is often regarded and measured as a single, homogeneous disorder and we have treated it as such in this paper. Importantly, diagnosis of depression requires only presence of one out of the two core symptoms: prolonged negative affect and the loss of interest in previously pleasurable activities (DSM-5; American Psychiatric Association, 2013). A growing amount of evidence is pointing to the heterogeneous nature of depression and the need to analyze specific symptoms of depression (Fried and Nesse, 2015). Our framework emphasizes the possibility that cognitive control deficits in depression emerge from different causes. This allows for a study of how potential different causes of cognitive control deficits in depression can be related to different symptom clusters. For example, reward processing impairments, resulting in lowered levels of cognitive control allocation, can be related to anhedonic symptoms. In the same way, changes in efficacy estimates can produce control deficits, but they could be more related to the presence of negative affect.

The study of mechanisms through which cognitive control deficits in depression emerge is of great relevance for developing future cognitive treatments for depression. While there is no evidence for the effectiveness of cognitive training in healthy individuals (Simons et al., 2016), cognitive trainings in depressed individuals are showing considerable promise (Koster et al., 2017; Motter et al., 2016; Siegle et al., 2007). However, the mechanisms through which cognitive trainings improve depressive symptoms remain unknown. Moreover, studies on cognitive training in depression have not focused on motivation yet. Understanding the components of motivation that are affected by these trainings, as well as including incentives in the trainings, will allow for more precise and individualized treatments.

## Conclusions

Cognitive control deficits represent an important vulnerability factor for depression and play a central role in cognitive impairments in depression. We propose a framework of cognitive control in depression in which these deficits emerge as a consequence of changes in the decision-making process about control allocation. We argue that alterations in core components of motivated behavior – reward anticipation, effort costs, and estimates of environment controllability – are crucial mechanisms contributing to dysfunctional cognitive control in depression. This view provides a more mechanistic understanding of cognitive control in depression, offers a better computational and neural understanding of the deficits, and connects cognitive research on depression with other fields of study such as motivation and agency. We believe that our framework can guide future research on mechanisms underlying depressive cognition which can result in improving treatments for depression.

## Acknowledgements

This work was supported by the Special Research Fund (BOF) of Ghent University [grant number 01D02415 awarded to IG], by a Center of Biomedical Research Excellence grant from the National Institute of General Medical Sciences [grant number P20GM103645 awarded to AS], by the starting grant of the European Research Council (ERC) under the Horizon 2020 framework [grant number 636110 awarded to RK], and by the Concerted Research Action Grant of Ghent University [grant number BOF16/GOA/017 awarded to EHWK], and a grant from the John Templeton Foundation [awarded to SM; The opinions expressed in this publication are those of the authors and do not necessarily reflect the views of the John Templeton Foundation].

## Supplementary Materials

### Simulations of control allocation

In order to simulate the effects of motivational variables on control allocation we generated behavior from an agent that performs cognitive control tasks^2^ (Musslick et al., 2015). In these simulations, each trial is described as an interaction between the control system and the task environment. The agent specifies the optimal control signal based on an internal representation that is an estimate of the next trial (inferred state *Ŝ*). The agent implements the specified control signal while performing the corresponding task in the actual task environment (actual state *S*). After each trial, the agent updates the inferred state for the next trial based on its observation of the current trial.

To generate response probabilities and reaction times we used a drift diffusion model (DDM, Bogacz et al., 2006; Ratcliff, 1978) that accumulates evidence towards one of two responses in a Stroop task (e.g. one response indicating the color green and the other response indicating the color red). The DDM simulates performance on a task as the accumulation of evidence about the stimulus towards one of two responses. The drift rate determines whether accumulated evidence leads to either the lower or the upper threshold (e.g. leading to either the left or right response button) and its magnitude determines how fast evidence is accumulated. The threshold determines the amount of evidence required to indicate a response, regulating the speed-accuracy tradeoff: higher thresholds lead to higher likelihood of reaching the correct response at the expense of slower reaction times (Ratcliff, 1985, 1978). We assume that the rate of accumulation toward one of the two response boundaries is governed by an automatic component and a controlled component

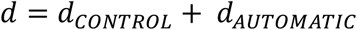

where the automatic component reflects automatic processing of each of the color and word of the stimulus that is unaffected by control:

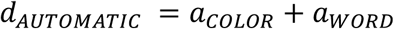

The absolute magnitude of the color-response association *a*_*COLOR*_, as well as the magnitude of the word-response association *a*_*WORD*_ depends on the strength of the association of each stimulus feature with a given response and its sign depends on the response (e.g. *a_COLOR_* < 0 if the response is associated with the left button, *a_COLOR_* > 0 if response is associated with the right button). Thus, for congruent trials *a*_*COLOR*_, and *a*_*WORD*_ have the same sign, whereas the opposite sign for incongruent (conflict) trials. The controlled component of the drift rate is the sum of the two stimulus values, each weighted by the intensity of the corresponding control signal, one for processing the color dimension of the stimulus *u_COLOR_* and one for processing the word dimension of the stimulus *u_WORD_*:

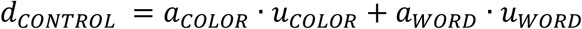

Thus, each control signal biases processing towards one of the stimulus dimensions. As a result, higher control signal intensity for processing the color dimension of the stimulus improves performance – speeds responses and lowers error rates – for the Stroop task. Mean reaction times (RTs) and response probabilities for a given parameterization of drift rate on trial *t* are derived from an analytical solution to the DDM (Navarro and Fuss, 2009).

In order to specify the optimal set of control signals *U* = {*u*_*COLOR*_, *u*_*WORD*_ } on a given trial, the model estimates the expected value for each configuration of control signal intensities based on its internal model of the next trial *Ŝ* = {*â*_*COLOR*_, *a*_*WORD*_ }. This is done by weighting the expected reward for an outcome against the cost associated with the chosen control signal configuration:

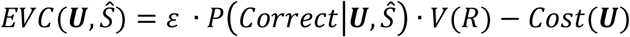

Where ε corresponds to the expected efficacy of exerting cognitive control, *P(Correct|**U**, Ŝ)* corresponds to the probability of reaching the decision threshold for the correct response and *V(R)* corresponds to the subjective value of responding correctly. Here, the subjective value *V(R) = v · R* corresponds to the amount of reward offered for a correct response *R* weighted by the model’s sensitivity to the reward *v*. The total cost of cognitive control *Cost(C)* is computed as the sum of the costs for each control signal,

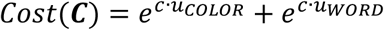

where the cost for each control signal is an exponential function of the intensity of the control signal, scaled by the cost parameter *c*. The model selects the control signal configuration with the maximum EVC within the inferred next trial *Ŝ*, out of all the configurations under consideration:

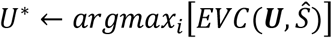

Performance in the actual state *S* is determined by the influence of the chosen control signals on the true parameters of the DDM (e.g., by adjusting the drift rate).

### Task environments

#### Stroop Paradigm

To illustrate effects of distractor interference in the EVC model, we simulated behavior on a Stroop task. In this task, the model is presented with a two-dimensional stimulus, one dimension representing an ink color and another dimension representing a color word. On each trial, the EVC model is required to indicate the response associated with the ink color. The trial sequence encompassed 200 trials, half of which were response congruent and half of which were response incongruent. To simulate congruent trials, we set *a*_*COLOR*_ = 0.1, *a*_*WORD*_ = 0.33 such that both stimuli dimension promote the same response. On incongruent trials, we set *a*_*COLOR*_ = 0.1, *a*_*WORD*_ = −0.33 such that the word dimension is associated with a different response than the color dimension. Note that the word-response association has a higher absolute magnitude than the color-response association, reflecting the assumption that word reading is a more automatic process than color naming. To simulate an expected mixture of congruent and incongruent trials, we parameterized the expected state as the average of the two trial conditions, *Ŝ* = {*â*_*COLOR*_ = 0.1, *a*_*WORD*_ = 0}. Control was implemented in the form of two control signals, one for processing the color dimension and one for processing the word dimension. The range of control signal intensities as varied from 0 to 4 in steps of 0.02 and the reward received for a correct response was set to *R = 40*. DDM parameters were set as follows: starting point = 0.0, noise coefficient = 1.0, non-decision time = 0.25s and threshold = 1.5. To demonstrate how control-demanding behavior is affected by changes in the decision-making process about control allocation we systematically varied the control cost parameter *c* from 0.5 to 1.5 in steps of 0.1, the reward sensitivity *v* from 0.5 to 1.5 in steps of 0.1 and the expected control efficacy *E* from 0.1 to 1.0 in steps of 0.1 across simulations. Note that we varied only one parameter at a time while holding the other parameters constant at *c* = 1, *v* = 1, *ε* = 1. For each parameter setting, we assessed the mean amount of control that the model allocated for each trial type, as well as the mean error rate.

#### COGED Paradigm

To demonstrate the effects of changing model parameters on demand-avoidance, we simulated behavior in the cognitive effort discounting (COGED) experiment described by Westbrook & Braver (2015). In this paradigm, subjects can choose on each trial whether they want to perform a baseline low-demand task for a low reward or a higher-demand alternative task for a higher reward. The amount of reward offered for the baseline task is adjusted to identify the point of indifference, that is, the reward at which subjects are indifferent between performing the low-demand baseline task and performing the high-demand task. To simulate this paradigm, we modeled both tasks as different types of trials that the model can choose between. Each trial encompassed a stimulus with a color dimension that mapped to one of two responses with *a_COLOR_* > 0. However, unlike in the Stroop task described above there was no word dimension, *a*_*WORD*_ = 0. The difficulty of the high-demand task was manipulated across experiment blocks by varying the color-response association *a*_*COLOR*_ from 0.2 to 0.4 in steps of 0.0667 and the difficulty of the baseline task was fixed to *a*_*COLOR*_ = 1 (higher color-response associations may reflect higher saturation values for a color patch). For each set of simulations, we fixed the reward for the high-demand task to (*R* = 200) while steadily increasing the amount of reward offered for the low-demand task in steps of 1, beginning from an initial reward value of *R* = 1. On each trial, the EVC agent determined the highest EVC separately for each task and chose the task with the highest predicted EVC. We then assessed the amount of reward offered for the low-demand task for which the model would be indifferent between performing the low-demand task and the (more rewarding) high-demand task and normalized this value by the amount of reward offered for the high-demand task. Following the notation by Westbrook & Braver (2015), we refer to this reward value as the subjective value of completing the high-demand task. For instance, if the model would switch to performing the low-demand task at an offered reward of 120 then the (discounted) subjective value of the high-demand task would be 120. The range of control signal intensities was varied from 0 to 6 in steps of 0.05 and DDM parameters were set as follows: starting point = 0.0, noise coefficient = 1, non-decision time = 0.25s and threshold = 1.5. To demonstrate how control-demanding behavior is affected by changes in the decision-making process about control allocation, we systematically varied the control cost parameter *c* from 0.5 to 1 to 1.5, the reward sensitivity *v* from 0.5 to 1 to 1.5 and the expected control efficacy *E* from 0.1 to 0.5 to 1.0 across simulations. Note that we varied only one parameter at a time while holding the other parameters constant at *c* = 1, *v* = 1, *ε* = 1. For each parameter setting, we assessed the subjective value as a function of the task difficulty of the high-demand task.

## Footnotes

1 Throughout this paper we use the term depression to denote the Major Depressive Disorder.

2 The source code for all simulations is available at https://github.com/musslick/EVCDepression

